# A Primary Central Source Determines Perturbation-Evoked N1 Amplitudes but not Latencies in Younger Adults

**DOI:** 10.64898/2026.01.23.701364

**Authors:** Janna Protzak, Jasmine L. Mirdamadi, Michael R. Borich, Lena H. Ting

## Abstract

The balance perturbation-evoked N1 potential is a reliable cortical response during reactive balance control that is correlated to a variety of cognitive and motor functions. Although the supplementary motor area (SMA) has been identified as the primary source of the N1, it is less understood whether other brain regions contribute to N1 recorded at the scalp. We used source localization on electroencephalography (EEG) data from 25 younger adults recorded during backward whole-body perturbations during stance. We identified the sources that contribute to channel-based N1 recordings and quantified their impact on N1 amplitude and latency. In younger adults, N1 amplitudes can be explained by one single source in a central midline cortical region covering the SMA. When reconstructing N1 signals using backprojections with one versus all independent components (IC) identified as brain sources there was no difference in peak amplitudes and a small but significant difference in N1 peak latencies. Parallel brain sources thus deflect the time course of the N1, but not its magnitude. Brain areas associated with IC’s contributing to the shift in N1 latency varied between participants. Our results emphasize the dominant influence of central cortical areas on the N1 response, informing hypothesizes regarding the nature of the signal and its functional role. Importantly, the extent and location of other cortical structures that influence N1 timing, such as parietal cortex areas and the anterior cingulate cortex, may further elucidate cortical contributions to balance. These markers could be crucial for the early detection of balance problems in clinical populations.

**NEW & NOTEWORTHY:** We demonstrate that channel-level amplitudes of the balance perturbation-evoked N1 in younger adults primarily reflect neural activity originating from cortical central midline regions, particularly the SMA. In contrast, contributions from parallel active brain regions evoked by balance perturbations are indicated by an influence on N1 peak latencies. Our findings imply that the perturbation-evoked N1, unlike other evoked potentials, is not a mixture of multiple neural sources in younger adults.

## INTRODUCTION

Electroencephalography (EEG) data offer precise temporal measures of cortical activity during balance control, with the disadvantage of poor spatial resolution. The signals measured at the electrode level reflect a combination of volume conducted brain sources, as well as non-brain activity (e.g., from muscles or external artifacts) (1,2). Alterations in cortical engagement patterns underlying reactive balance control may foreshadow balance impairments associated with aging or cortical pathologies [see (3,4) for overview articles]; therefore, a detailed analysis of the relative contributions of cortical structures to well-studied perturbation-evoked scalp potentials is important. The most prominent perturbation-evoked scalp potential is the N1, which is a large negative deflection recorded over central EEG electrodes 100 to 200 milliseconds after externally elicited perturbations to the body, e.g. support surface translations during stance [see (5,6) for overview articles]. While the N1 has previously been localized to the supplementary motor area [SMA, (7,8)], the extent to which other brain sources contribute to the N1 remains unclear.

Standing balance perturbations are usually salient events that move the whole body, triggering rapid balance-correcting actions to an abrupt external event. Although perturbations can cause multisensory signals, including vestibular and visual information, proprioception may play a more dominant role (9). However, the N1 is most likely not dependent on one modality, as it can be detected when participants have their eyes closed (10), and in a patient with bilateral labyrinthine loss (11) or during seated conditions (12,13). The N1 is further a highly reliable signal that is typically detected in every participant and on a single-trial basis with prominent amplitudes (8,14,15) that can scale with perturbation intensity (14,16,17). The characteristics of the exogenous physical perturbations alone are, however, insufficient to explain the observed variability in N1 characteristics. Typically, the amplitude is largest on the first trial of an experiment (8) smaller for predictable than unpredictable perturbation (18) and attenuate with repetitions (8,19,20), suggesting modulatory influence of top-down processes.

The involvement of the SMA is an indication that the N1 may reflects both bottom-up and top-down neural processes due to its connections to prefrontal and cingulate areas and direct projections to the spinal cord (21,22). Connections between the SMA and the basal ganglia have further put this brain region in the focus of research centered on the motor deficits in Parkinson disease (23). Functionally, the SMA has been associated with initiation of internally driven movements or changes in movement plans [see (22) for an overview article], e.g. when an external cue signals that an already initiated movement has to be altered. However, previous studies that have localized the main source of the N1 to Brodman area 6 or specifically to the SMA (7,8) lack a detailed assessment of how much other brain areas influence the N1 potential recorded at channel level. Consequently, it is not known whether N1 reflects a mixture of multiple parallel active neural sources or a signal that can be primarily attributed to the SMA, complicating the functional interpretation.

Here, we reanalyzed data from a sample of younger adults and used independent component analysis (ICA) to describe brain sources contributing to the perturbation-evoked N1, recorded at channel level. We hypothesize the large negative deflection measured at central scalp locations following perturbations is a signal predominately reflecting SMA activity. We predicted that the time course of activation at the central midline channel with the most prominent N1 would be highly correlated with the time course of activation of the back-projected N1 using the *primary contributing IC* and that this IC would be predominately located in brain regions covering the SMA. We further explored effects of concurrent activity of other identified sources on the amplitude and timing of the N1 response.

## MATERIALS AND METHODS

### Participants

Data from 25 healthy younger adults (12 female, mean age: 24 years, SD: 3.9 years) were reanalyzed for the present report. Initial results from this sample have been reported in Mirdamadi et al. 2024 (20) (n=15) and Mirdamadi et al. 2025 (15)(n=10). Written informed consent was given by all participants and the study protocol was approved by the Emory University Institutional Review Board.

### Experimental task and setup

For the current analyses, we extracted data from a whole-body motion-perception task in which participants were asked to judge whether two consecutive backward perturbations were in the same or different direction. We only considered the first perturbation of each perturbation pair for our analysis, which was always a straight backward perturbation. These perturbations had a consistent characteristic (7.5*cm* displacement, 15*cm/s* velocity, 0.1*m/s*^2^ peak acceleration), but were delivered at unpredictable time intervals to minimize predictability and adaptation.

### EEG recordings and preprocessing

EEG data were recorded continuously with a 64-channel EEG system (ActiCHAMP, Brain Products GmbH, Gilching, Germany) at a sampling rate of 1000Hz. All electrodes were active amplified and arranged according to the extended international 10%-system (24). A ground electrode was positioned at location AFz and the reference channel at Fz. The raw EEG data sets were preprocessed using custom Matlab scripts and the EEGLAB(25)-based BeMoBIL pipeline for mobile brain/body imaging data (26). The raw data sets were loaded into the EEGLAB-format and the timing of the perturbation triggers were corrected based on the data of an accelerometer, which was attached to the platform and recorded synchronously with the EEG data through an onboard auxiliary channel of the EEG amplifier.

Basic preprocessing of the EEG data included the assignment of standard coordinates to each electrode label and the removal of line noise (27,28). Noisy channels were identified using the *clean_artifacts* function of the EEGLAB *clean_rawdata* plugin with a [0.25 0.75] transition band high pass filter, a channel correlation criterion of 0.7 as a minimal correlation coefficient and a maximal time window of 300ms that had to be exceeded to flag a channel as bad. Since this function uses a random data sample, which can produce unstable results, the channel rejection procedure was repeated twenty times. Only channels that were flagged in more than a third of all iterations were rejected. Subsequently the rejected channels were reconstructed using spherical interpolation and all channels were re-referenced to a common average, and the data from the original reference channel Fz was reconstructed.

For source localization, preprocessed data sets were rank-reduced by the number of interpolated channels, high-pass filtered with a cutoff frequency of 1.5 Hz (29) and epoched to a time window of 2s before to 2s after each perturbation. Subsequently each data set was decomposed using adaptive mixture independent component analysis (AMICA) (30,31). 10 iterations of the AMICA build-in functionality that removes data samples with a log-likelihood lower than 3 SD were used to improve data quality (31). IC dipole locations were fitted using the EEGLAB DIPFIT toolbox and a standard boundary element method (BEM) head model (32).

The obtained AMICA weights and dipole models were extracted and projected to the initial basic preprocessed continuous data sets with a high-pass filter of 0.5 Hz (lower-edge). In a final data cleaning step, ICs were classified as representing brain activity or as artifact IC’s based on the following steps. First, all ICs were rejected if a) the residual variance exceeded 15 percent, b) the IClabel toolbox (33) (default classifier) classified the IC as representing eye activity, line or channel noise as the category with the highest probability c) if the frequency spectrum had a positive slope (≥ 0.31) which can be indicative of muscle activity (34), or d) if at least one electrode exhibits a z-score > 5, based on the IC topography weights calculated and visualized with the TESA pluggin (35). All resulting ICs were visually checked based on the validity of their dipole location and corresponding IC scalp maps, power spectrum and the cross-frequency power–power coupling using the powpowcat toolbox (36). On average, 10 ICs (SD=2.6) were identified as brain ICs and backprojected to the channel level.

### EEG data analysis

The resulting data sets, containing the channel level data as well as the ICA weights, were lowpass-filtered with a pass-band edge of 30 Hz. Subsequently, each set was epoched into trials (−1 to 2 s relative to the perturbation onset) and baseline corrected to the pre-perturbation time window of −200 to −50 ms. Trials exceeding a threshold of ±100*µ*V were rejected. On average, 0.3 trials were rejected (SD= 0.6) and 61 trials (SD= 2.2) per participant entered the analysis. The cleaned and epoched data sets were used to characterize the reconstructed channel N1 potential for the *primary IC* contributing to the N1 as well as for the backprojection of *all brain ICs*. The analysis pipeline described in the following is depicted for two exemplary participants in Fig. 1.

**Figure 1.**
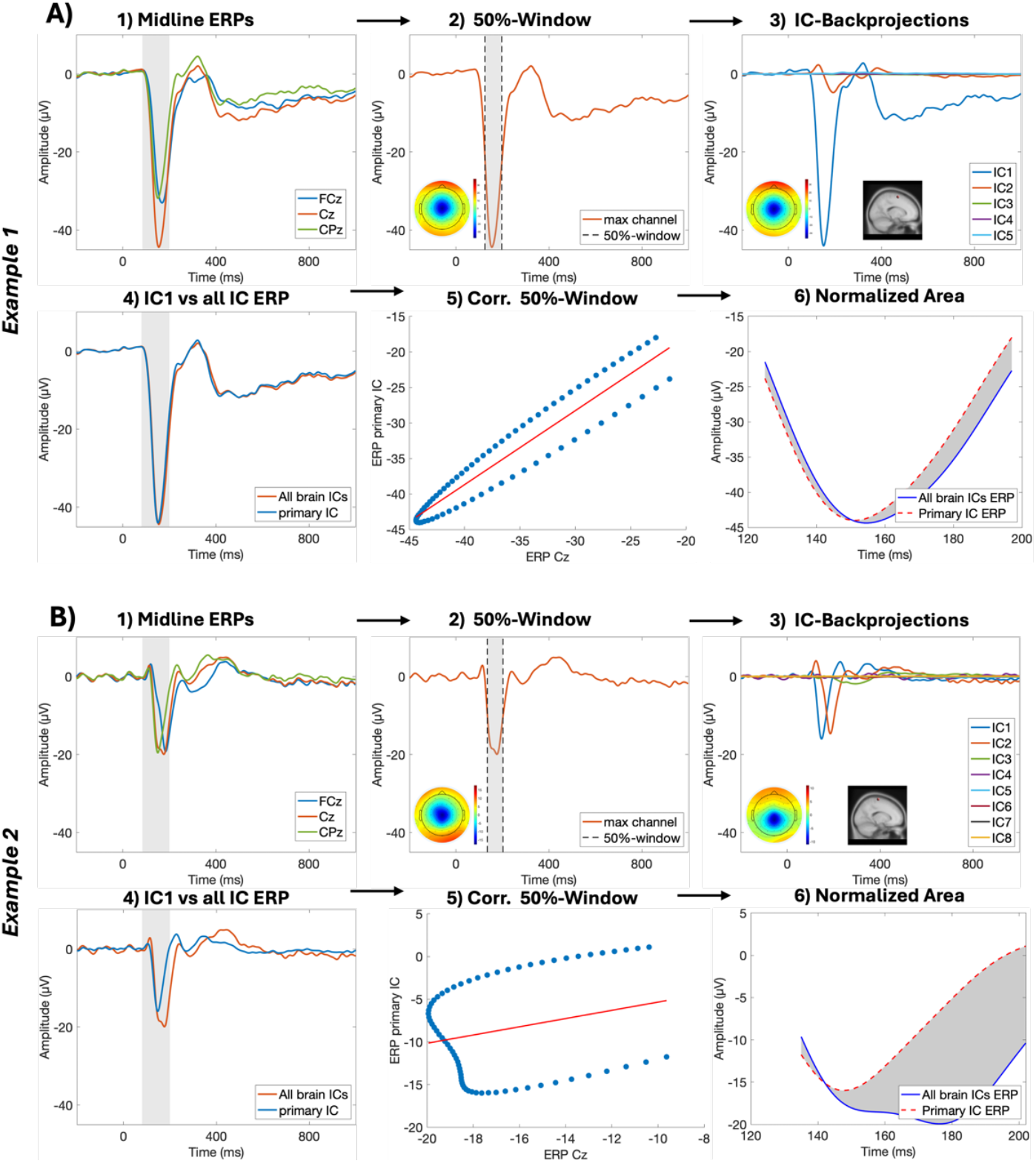
Analysis pipeline for two exemplary participants: upper part A) for a typical participant and bottom part B) for one participant with the highest discrepancy between *primary N1* and *all brain IC*-backprojections. 1) Time-activation course for midline electrodes FCz, Cz and CPz and the N1-peak extraction window in gray; 2) the individual 50% data extraction window for the N1 for the channel with largest negativity within the N1-time window and the topography for that time window; 3) the backprojection of each brain IC to the individual channel identified in step 1, i.e. the contribution of each IC to the channel and the IC scalp map and location of the *primary IC*; 4) the total backprojection of *all brain ICs* (red) and the *primary N1 IC* (blue) to the individual N1-channel, with the N1 time window in gray; 5) the correlation between all data points within the 50%window between the traces of the backprojection of *all brain ICs* in red and for the primary N1-IC; 6) the area difference between the curves of the backprojection of *all brain ICs* in red (dashed line) and for the *primary N1 IC* in blue in the 50% window.

For the electrode selection, we plotted the time-activation course at the three midline electrodes FCz, Cz and CPz from the backprojection of *all brain ICs* to channel level and determined the electrode with the highest negativity in the N1 time window between 80 to 200ms after perturbation onset (Fig. 1.1). We chose the most commonly reported N1 location, Cz, plus the two neighboring midline electrodes, FCz and CPz as our area of interest to account for individual differences in functional neurotomy and the inherent spatial imprecision of EEG recordings. Each individual’s N1 time window for analysis was specified as the full width of the negative deflection at 50% of the peak amplitude, referred to as the 50%window (Fig. 1.2). For each participant, the IC with the highest negativity within this time window was selected as the *primary N1 IC* with the highest contribution to the potential measured at the N1-channel (Fig. 1.3).

**Figure 2.**
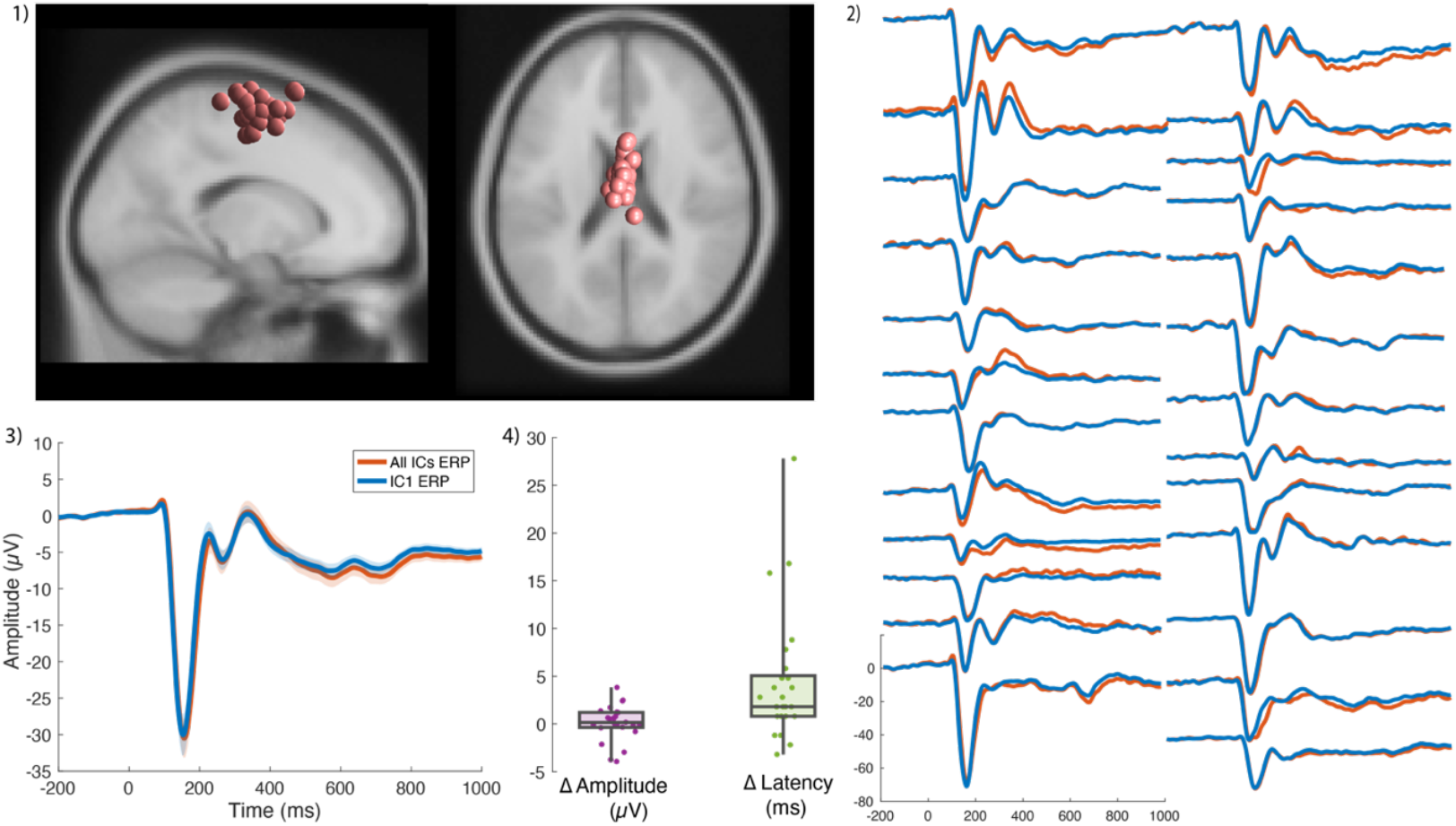
1) MNI coordinates for the individual *primary N1IC*; 2) single subject ERP traces with *all IC*-backprojections in purple and *primary N1 IC*-backprojections in green (x-axis: ms, y-axis: *µ*V); 3) group-average ERP with the shaded region showing the standard error of the mean; the *all brain IC* backprojection is shown in red and *primary N1 IC-*backprojection in blue, 4) single subject differences (Δ) in amplitude in purple on the left and in green for latencies on the right.

**Figure 3.**
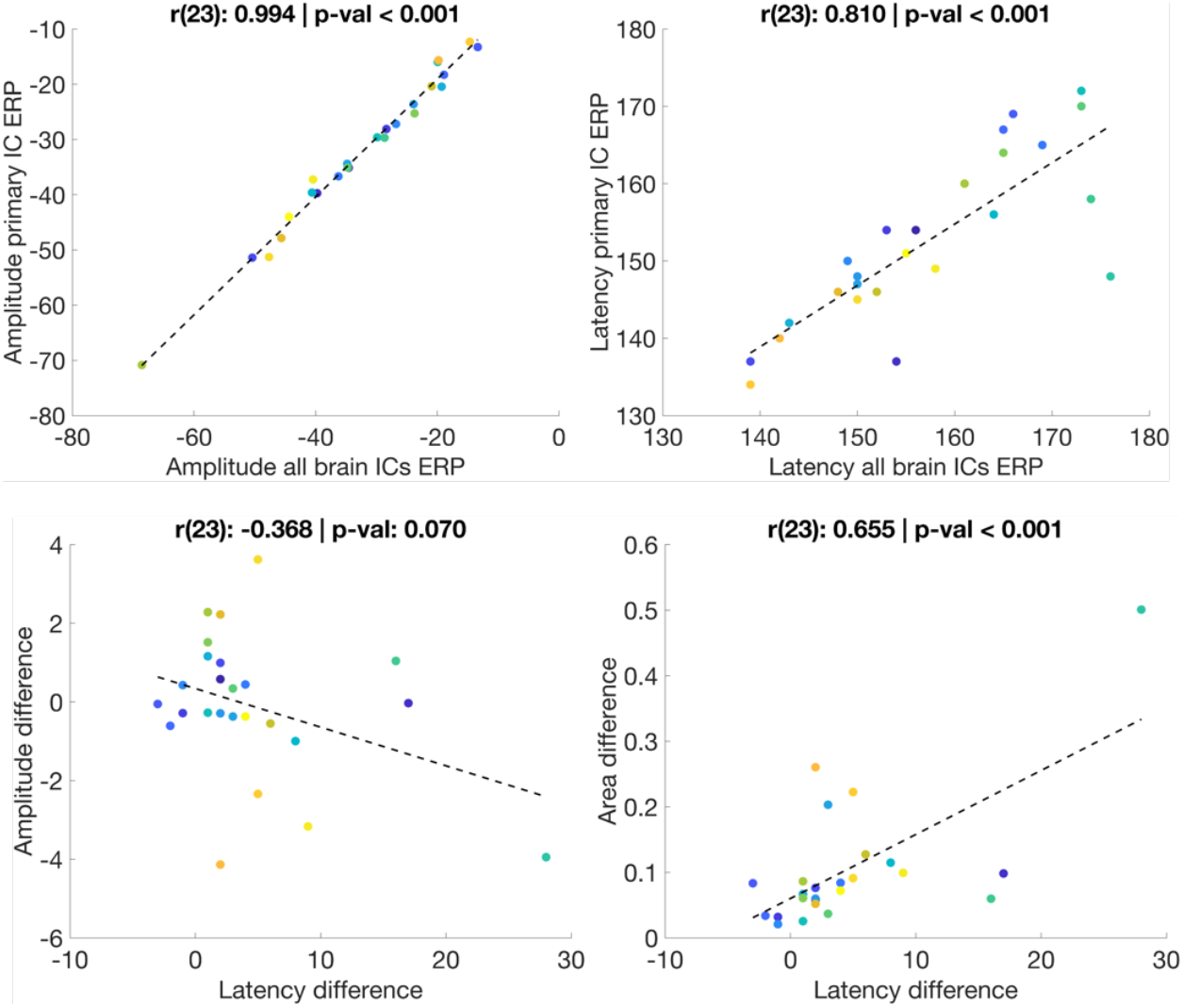
Correlations between the peak amplitudes (in *µ*V) of the *all brain ICs* backprojection and the *primary N1 IC* backprojection (upper left); between the peak latencies (in *ms*) of the *all brain ICs* backprojection and the *primary N1 IC* backprojection (upper right); between the differences in peak latencies (in *ms* on the x-axis) and differences in peak amplitude (y-axis in *µ*V, lower left); between the latency differences (in *ms* on the x-axis) and area differences (y-axis, lower right)

Peak amplitudes and latency measures were extracted for the backprojection of *all brain ICs* and for the backprojection of the *primary N1 IC* (Fig. 1.4). The MNI coordinates of each individual *primary N1 IC* were associated with a brain region using the *BioImage Suite Web* application (https://bioimagesuiteweb.github.io/webapp/mni2tal.html). Approximate brain regions of the second highest contributing ICs are reported as group means. The association of the activation time series of the N1 for both backprojections - the *primary N1 IC* backprojection and the *all brain IC* backprojection - within the 50%-window were analyzed using Pearson correlation (Fig. 1.5). Differences between the curves were further quantified as the area between the curves of the N1-channel for the *all brain IC* backprojection and the *primary N1 IC*-backprojection. We integrated the absolute difference between both curves within the 50%-window using the matlab trapz.m function. The area difference is reported as a percentage of the area under curve of the N1 for the *all brain IC* backprojection (Fig. 1.6). The mean values and standard deviations of the correlations and area measures are reported to describe the participant-level data.

### Statistical analysis

For each participant, the peak amplitude and corresponding latency were extracted for statistical analysis using the N1-channel that was reconstructed using backprojections of a) *all brain ICs* and b) only the *primary N1 IC*. Differences in amplitudes and latencies were assessed with paired t-tests.

Pearson’s correlation coefficients were used to assess associations between N1 latencies and amplitudes of both backprojections. To further explore potential association of the significant differences in peak latencies, post-hoc correlations were calculated with the differences in amplitudes and with the differences in area under the curve in the 50%window (expressed as the percentage change from the *all brain IC*s to the *primary N1 IC* back-projection).

## RESULTS

### Descriptive subject level results

This section summarizes the descriptive subject level results for the whole sample, which are depicted in Fig. 1 for two exemplary participants: for one participant with results representing the group mean and for the participants with the highest area difference between the two backprojections. All single subject primary ICs and ERP traces are shown in Fig. 2.1 and Fig. 2.2. The grand average ERPs are shown in Fig. 2.3.

Channel Cz was the central channel with the highest negativity in the N1 time window for 19 participants; for 5 participants it was channel CPz and for one participant it was channel FCz (see Fig. 1.1). For the individually determined N1-channel, the average time window spanning the full width of the N1 at 50% of the peak amplitude was 64.32*ms* (*SD* = 15.09) and ranged from 39ms to 103ms (see Fig. 1.2).

The within-participant correlations between the curves of the *primary N1 IC* and *all brain ICs* backprojection within the 50%window ranged from 0.24 to 1.00, with a mean of *r* = 0.90 (*SD* = 0.16), cf. Fig 1.5. The average area between both curves was 10.5% (*SD* = 10.19) of the total area under the N1 of the *all brain ICs* back-projection, with a range of 2.1% to 50.1% (cf. Fig. 1.6).

The *primary N1 IC* was typically localized within Brodmann Area 6, a brain region covering the medially located SMA as well as lateral premotor areas (Table 1 and Fig. 2.1). For 14 participants, the *primary N1 IC* was located within the central part of Brodmann Area 6 and thus in the part that has been associated with the SMA. For 9 participants, the *primary N1 IC* was located outside of defined brain areas, but near the central part of Brodmann Area 6 (e.g. within the central sulcus). For one participant, the *primary N1 IC* was located on the border of Brodmann Area 6 and Brodmann Area 24. The latter comprises the anterior cingulate cortex (ACC). For one participant, the *primary N1 IC* was located within Brodmann Area 4 which covers the primary motor cortex area.

**Table 1.** Mean MNI coordinates and associated brain areas for the individual *primary N1 ICs* as well as the cluster centroid. Parentheses indicate that the exact coordinates are outside defined areas, e.g. in the central sulcus. the nearest area is indicated within the brackets. BA 4: Brodmann Area 4: Primary Motor Cortex, BA6: Brodmann Area 6/SMA and premotor areas, BA 24: Brodmann Are 24/ anterior cingulate cortex.

| ID | MNI (x,y,z) | Area | ID | MNI (x,y,z) | Area |
| --- | --- | --- | --- | --- | --- |
| 1 | 5, 1, 55 | BA 6 | 14 | 1, -19, 72 | (BA 6) |
| 2 | -1, -3, 66 | BA 6 | 15 | 1, -12, 52 | (BA 24/ BA 6) |
| 3 | -4, -7, 60 | BA 6 | 16 | -1, -15, 61 | BA 6 |
| 4 | 0, -7, 71 | (BA 6) | 17 | 6, -29, 69 | BA 4 |
| 5 | -2, -17, 60 | BA 6 | 18 | -2, -15, 69 | BA 6 |
| 6 | 1, 2, 61 | BA 6 | 19 | 2, 11, 74 | (BA 6) |
| 7 | -2, -13, 75 | (BA 6) | 20 | -2, -5, 64 | BA 6 |
| 8 | -7, -15, 51 | BA 6 | 21 | 3, 11, 74 | (BA 6) |
| 9 | 3, -7, 63 | (BA 6) | 22 | 1, 6, 642 | (BA 6) |
| 10 | -5, -20, 74 | BA 6 | 23 | -2, -8, 65 | BA 6 |
| 11 | -2, -13, 63 | (BA 6) | 24 | 5, 2, 65 | BA 6 |
| 12 | -1, -11, 60 | BA 6 | 25 | 0, -3, 63 | (BA 6) |
| 13 | 2, -8, 57 | BA6 | mean | 0, -8, 64 | (SMA) |

For twelve participants the IC with the second highest negativity within the 50% time window was located in Brodmann Area 24 and 32, i.e brain regions including the ACC. For three participants, the second IC was located in Brodmann Area 23 and 31; brain regions that comprise the posterior cingulate cortex. For three participants, the second IC was located in Brodmann Area 6, a region that spans the SMA and premotor area. Other brain areas for the second IC included Brodmann Area 8 (frontal eye field) for two participants, Brodmann Area 5 (somatosensory association cortex) for two participants, Brodmann Area 18 (visual association area V2) for one participant, the caudate for one participant and Brodmann Area 7 (precuneus/ superior parietal lobule) for one participant. On average, the second IC peaked 24.2ms (*SD:* 22.4ms) after the first IC.

### Group Level results

The mean N1 peak amplitude of the *all brain IC* backprojection was −32*µ*V, (*SD* = 13) and −32*µ*V, (*SD* = 11) for the *primary N1 IC* backprojection (*t*(24)=−0.308, *p* = .761), with a mean difference of −0.11*µ*V, (*SD* = 1.82) (Fig. 2.3).

The mean N1 peak latency of the *all brain IC* backprojection was 156*ms*, (*SD* = 10) and 152*ms*, (*SD* = 10) for the *primary N1 IC* backprojection (*t*(24)= 3.37, *p* = .003), with a mean difference of 4.60*ms*, (*SD* = 6.83) (Fig. 2.4). The effect for latencies remained significant after correction for one outlier with a latency difference *>* 3*SD* away from the mean.

The correlation between the *all brain IC* backprojection and the *primary N1 IC* backprojection was *r*(23)= 0.99, *p <* .001 for the peak amplitudes and *r*(23)= 0.81, *p <* .001 for the corresponding latencies (see Fig. 3). The differences in latencies between both back-projections was not correlated with the differences in peak amplitudes (*r*(23)=−0.37, *p* = .070), but with the normalized area difference (*r*(23)= 0.65, *p <* .001, see Fig. 3). However, the correlation with the area difference was driven by one participant with a value that is more than 3 SD away from the mean. Following exclusion of this participant, the correlation was no longer significant (*r*(22)= 0.19 *p* = .380).

## DISCUSSION

We found that the balance-perturbation evoked N1 responses in younger adults reflect almost exclusively activity from a single source located in central brain areas corresponding to the SMA, while differences in the peak timing of the N1 indicate parallel activity from a larger cortical network active during reactive balance control. Although a dominant N1 source was identified for each participant, the source of the secondary parallel activity varied between participants. These differences in cortical engagement during whole-body perturbation could be crucial for our understanding of cortical involvement pattern underlying voluntary balance control, particularly for characterizing balance impairments.

### N1 amplitudes in younger adults can be explained by one source in or near the SMA

Our results demonstrate that the balance perturbation-evoked N1 in younger adults recorded at central scalp locations originates from a single source located in or near the SMA.

Importantly, we did not find evidence to suggest that other sources influence its magnitude. Using a relatively simple approach, we found one single IC for each participant that explains most of the N1 amplitude. We were able to show that the average N1 peak from the backprojection of one single IC per participant is nearly identical in shape and highly correlated with the N1 extracted from the backprojection of the complete brain IC solution. The sources of the identified ICs that contribute primarily to the channel ERP, as measured by the reconstruction of all brain ICs, form a tight cluster around central brain regions associated with the SMA with relatively little spread. While EEG data can only provide rough estimates of brain source locations, the large N1 amplitude signals presumably led to better signal-to-noise ratios and a relatively stable source localization. In our study, this resulted in a tight group-level IC cluster with a centroid near the central sulcus close to Brodmann area 6. Based on the medial location of the cluster and the MNI coordinates of the individual ICs, we argue that the origin is within the medial SMA part and not in the lateral premotor areas of Brodmann area 6. Despite the inherent limitations in spatial resolution of the EEG data and the absence of co-registration with the individual MRI data in the present study, we observe prevalent sources within the approximate SMA region, which replicates prior results.

Therefore, we propose, that the whole-body balance N1 can provide a direct readout of evoked activity of brain structures covering the SMA during balance perturbations. Previous studies have demonstrated that the balance perturbation N1 can be reliably assessed in a single trial using a minimal and simple configuration consisting of six trials, three channels, and only basic preprocessing, such as high and low pass filtering (10). Our present findings add that the balance perturbation-evoked N1 in younger adults is nearly exclusively generated in one central brain region. Given the negligible influence from other sources on the N1 magnitude, it can be assumed that even with setups that contain fewer electrodes and minimal preprocessing, activity from a brain region comprising the SMA will be predominantly captured.

### Parallel activity might be masked by the dominant central N1

The high N1 amplitudes found in the present study, with a group mean around −30mV and up to −69mV for one participant, replicate previous findings (5), and may mask parallel activity at smaller scales. The very large N1 magnitudes are a distinctive attribute of balance-perturbation evoked cortical responses, as ERPs typically reflect propagated multi-source activity at lower magnitudes that often only manifests in multi-trial averages. For example, in studies using low to very intense auditory stimuli, average amplitudes have been reported to range from −1 to −15mV [see (37) for an overview article]. The observed differences in magnitude of the balance N1 compared to those evoked by other sensory stimuli are challenging to attribute to factors other than the nature of the stimulus itself, including potential differences in hardware or preprocessing methodologies (e.g., the EEG reference scheme). Whole-body perturbations are triggered by large proprioceptive or cutaneous bursts of activity and trigger brainstem-mediated involuntary balance-correcting responses that counteract the net direction of body motion. Our prior work show that the N1 may lie along a parallel transcortical pathway that evokes later cortically-mediated muscle activity (38,39) that may include strategy changes such as stepping responses (16,40). Unlike auditory stimuli, balance perturbations have inherent ecological relevance failure to respond appropriately can lead to a fall. The relatively large N1 response may thus indicate the negative consequences of the stimuli, but may also mask the activation of parallel cortical network activity (41) measured at much lower amplitudes at the scalp level.

### Additional sources characterize the timing of the N1

Our data support the previously reported finding that the channel-level, perturbation-evoked N1 in younger adults primarily arises from event-related activity in one central brain region and that additional sources influence N1 peak latencies. Although the practical implications of an average peak difference of less than 5ms might be questionable, this shift indicates that parallel sources are active during balance perturbations. Activity from these sources may not be large enough to deflect the N1 magnitude, but they are potentially within common amplitude ranges and can significantly affect the N1 timing. Especially since the functional role of the N1 is still under debate (5,42), contributions of additional sources to successful balance responses may be equally important. We scanned the remaining ICs with the second highest contribution to the N1, but we were unable to identify a single, spatially defined activity cluster. Sources with pronounced event-related activity within the N1 time range were distributed from frontal to parietal regions. Several participants in our sample exhibited significant negativities in two sources within the N1 time-window (for an example see Fig. 1.3). The participant with the lowest correlation between both curves and the largest amplitude and latency deviations revealed one large negative deflection around 150ms after perturbation onset in or near the SMA with the typical N1 timing and location and one smaller negativity around 30ms later in the anterior cingulate cortex (ACC). These findings may indicate parallel activity or reciprocal connections between the two structures during reactive balance control. As the ACC is a structure associated with conflict monitoring and error processing, one might speculate, that this structure is only triggered in participant with balance challenges and trials that require additional cortical involvement beyond regions around the SMA, and an initial brain stem modulated balance response. However, it is important to note that the analyzed data for this report were part of a complex perception task. Participants were asked to determine whether the direction of a subsequent perturbation was in a different direction than the straight-backward perturbations analyzed here. Therefore, it is necessary to determine whether the reported secondary sources are associated with the balance response or potential secondary task demands.

### What insights can the source contribution to the N1 reveal about its functional role?

Individual source profiles of parallel activities in combination with the behavioral measure of the balance response may be key for our understanding of the functional role of the perturbation-evoked N1. It is intriguing to argue that the N1 is a top-down modulated response evoked by the large proprioceptive input of a whole-body perturbation. However, based on the functionalities contributed to the SMA (43–46), the N1 could also be interpreted as a motor preparation process for a voluntary response or even as a ramped-up readiness-potential that builds-up in parallel to the first brainstem modulated motor response leading up to the onset or inhibition of an additional slower cortical mediated motor responses. Following this line of reasoning, the N1 could be a sensorimotor threshold correlate (44,47) for the time-sensitive motor preparation for a sudden change of the environment. Importantly, this would indicate that the N1 is not necessarily related to a motor output and thus might not directly related to behavioral measures as the perceptual and central processing could also result in the inhibition of a cortical modulated response. Examining populations with balance problems may provide further insight into this debate. It has been hypothesized that changes in activation patterns, such as more widespread cortical activation in older adults compared to younger adults performing the same task, are compensatory for age-related declines (48,49). Similarly, shifts in the contributions of different sources to the balance-evoked response may clarify their functional relevance. The composition of sources contributing to the N1 might be a candidate marker for the identification of early balance problems in aging and clinical populations, as the timely involvement of additional cortical structures might reflect a switch from automatic to balance responses requiring cortical compensation.

## DATA AVAILABILITY

The data sets will made openly available before publication.

## Acknowledgements

The authors acknowledge Scott Boebinger, Kennedy Kerr, Rish Rashtogi, Alex Poorman, Gaetan Munter, Kendra Jones, Andrew Monaghan and Patrick Knox for assistance with data collection.

## AUTHOR CONTRIBUTIONS

JM, LT and MB conceived and designed research; JM performed experiments; JP and JLM analyzed data; JP, JM, LT and MB interpreted results of experiments; JP prepared figures; JP drafted manuscript; JP, JM, LT and MB edited and revised manuscript; JP, JM, LT and MB approved final version of manuscript.

## REFERENCES

1. Gramann K, Ferris DP, Gwin J, Makeig S. Imaging natural cognition in action. Int J Psychophysiol. 2014 Jan;91(1):22–9.

2. Makeig S, Gramann K, Jung TP, Sejnowski TJ, Poizner H. Linking brain, mind and behavior. Int J Psychophysiol. 2009 Aug;73(2):95–100.

3. Monaghan AS, Takla T, Ofori E, Peterson DS, Wu W, Fritz NE, et al. The Neural Contributions to Reactive Balance Control: A Scoping Review of EEG, fNIRS, MRI, and PET Studies. Brain Sci. 2025 Dec 13;15(12):1330.

4. Huang HJ, Ferris DP. Non-invasive brain imaging to advance the understanding of human balance. Curr Opin Biomed Eng. 2023 Dec;28:100505.

5. Varghese JP, McIlroy RE, Barnett-Cowan M. Perturbation-evoked potentials: Significance and application in balance control research. Neurosci Biobehav Rev. 2017 Dec;83:267–80.

6. Purohit R, Bhatt T. Mobile Brain Imaging to Examine Task-Related Cortical Correlates of Reactive Balance: A Systematic Review. Brain Sci. 2022 Nov 2;12(11):1487.

7. Marlin A, Mochizuki G, Staines WR, McIlroy WE. Localizing evoked cortical activity associated with balance reactions: does the anterior cingulate play a role? J Neurophysiol. 2014 June 15;111(12):2634–43.

8. Mierau A, Hülsdünker T, Strüder HK. Changes in cortical activity associated with adaptive behavior during repeated balance perturbation of unpredictable timing. Front Behav Neurosci [Internet]. 2015 Oct 14 [cited 2025 Dec 18];9. Available from: http://journal.frontiersin.org/Article/10.3389/fnbeh.2015.00272/abstract

9. Dietz V, Quintern J, Berger W. Cerebral evoked potentials associated with the compensatory reactions following stance and gait perturbation. Neurosci Lett. 1984 Sept;50(1–3):181–6.

10. Mirdamadi JL, Poorman A, Munter G, Jones K, Ting LH, Borich MR, et al. Excellent test-retest reliability of perturbation-evoked cortical responses supports feasibility of the balance N1 as a clinical biomarker. J Neurophysiol. 2025 Feb 24;jn.00583.2024.

11. Dietz V, Quintern J, Berger W, Schenck E. Cerebral potentials and leg muscle e.m.g. responses associated with stance perturbation. Exp Brain Res. 1985;57(2):348–54.

12. Staines WR, McIlroy WE, Brooke JD. Cortical representation of whole-body movement is modulated by proprioceptive discharge in humans. Exp Brain Res. 2001 May;138(2):235–42.

13. Mochizuki G, Sibley KM, Cheung HJ, Camilleri JM, McIlroy WE. Generalizability of perturbation-evoked cortical potentials: Independence from sensory, motor and overall postural state. Neurosci Lett. 2009 Feb;451(1):40–4.

14. Payne AM, Hajcak G, Ting LH. Dissociation of muscle and cortical response scaling to balance perturbation acceleration. J Neurophysiol. 2019 01;121(3):867–80.

15. Mirdamadi JL, Poorman A, Munter G, Jones K, Ting LH, Borich MR, et al. Excellent test-retest reliability of perturbation-evoked cortical responses supports feasibility of the balance N1 as a clinical biomarker. J Neurophysiol. 2025 Feb 24;jn.00583.2024.

16. Solis-Escalante T, Stokkermans M, Cohen MX, Weerdesteyn V. Cortical responses to whole-body balance perturbations index perturbation magnitude and predict reactive stepping behavior. Eur J Neurosci. 2021 Dec;54(12):8120–38.

17. Goel R, Ozdemir RA, Nakagome S, Contreras-Vidal JL, Paloski WH, Parikh PJ. Effects of speed and direction of perturbation on electroencephalographic and balance responses. Exp Brain Res. 2018 July;236(7):2073–83.

18. Adkin AL, Quant S, Maki BE, McIlroy WE. Cortical responses associated with predictable and unpredictable compensatory balance reactions. Exp Brain Res. 2006 June;172(1):85–93.

19. Payne AM, Hajcak G, Ting LH. Dissociation of muscle and cortical response scaling to balance perturbation acceleration. J Neurophysiol. 2019 Mar 1;121(3):867–80.

20. Mirdamadi JL, Ting LH, Borich MR. Distinct Cortical Correlates of Perception and Motor Function in Balance Control. J Neurosci. 2024 Apr 10;44(15):e1520232024.

21. Dum R, Strick P. The origin of corticospinal projections from the premotor areas in the frontal lobe. J Neurosci. 1991 Mar 1;11(3):667–89.

22. Nachev P, Kennard C, Husain M. Functional role of the supplementary and pre-supplementary motor areas. Nat Rev Neurosci. 2008 Nov;9(11):856–69.

23. Jahanshahi M, Jenkins IH, Brown RG, Marsden CD, Passingham RE, Brooks DJ. Self-initiated versus externally triggered movements: I. An investigation using measurement of regional cerebral blood flow with PET and movement-related potentials in normal and Parkinson’s disease subjects. Brain. 1995;118(4):913–33.

24. Chatrian GE, Lettich E, Nelson PL. Ten Percent Electrode System for Topographic Studies of Spontaneous and Evoked EEG Activities. Am J EEG Technol. 1985 June;25(2):83–92.

25. Delorme A, Makeig S. EEGLAB: an open source toolbox for analysis of single-trial EEG dynamics including independent component analysis. J Neurosci Methods. 2004 Mar;134(1):9–21.

26. Klug M, Jeung S, Wunderlich A, Gehrke L, Protzak J, Djebbara Z, et al. The BeMoBIL Pipeline for automated analyses of multimodal mobile brain and body imaging data [Internet]. 2022 [cited 2025 Mar 1]. Available from: http://biorxiv.org/lookup/doi/10.1101/2022.09.29.510051

27. De Cheveigné A. ZapLine: A simple and effective method to remove power line artifacts. NeuroImage. 2020 Feb;207:116356.

28. Klug M, Kloosterman NA. Zapline-plus: A Zapline extension for automatic and adaptive removal of frequency-specific noise artifacts in M/EEG. Hum Brain Mapp. 2022 June 15;43(9):2743–58.

29. Klug M, Gramann K. Identifying key factors for improving ICA-based decomposition of EEG data in mobile and stationary experiments. Eur J Neurosci. 2021 Dec;54(12):8406–20.

30. Palmer JA, Kreutz-Delgado K, Makeig S. AMICA: An adaptive mixture of independent component analyzers with shared components. San Diego: Swartz Center for Computatonal Neursoscience, University of California San Diego; 2012. Report No.: 35.

31. Hsu SH, Pion-Tonachini L, Palmer J, Miyakoshi M, Makeig S, Jung TP. Modeling brain dynamic state changes with adaptive mixture independent component analysis. NeuroImage. 2018 Dec;183:47–61.

32. Oostenveld R, Praamstra P, Stegeman DF, Van Oosterom A. Overlap of attention and movement-related activity in lateralized event-related brain potentials. Clin Neurophysiol. 2001 Mar;112(3):477–84.

33. Pion-Tonachini L, Kreutz-Delgado K, Makeig S. ICLabel: An automated electroencephalographic independent component classifier, dataset, and website. NeuroImage. 2019 Sept;198:181–97.

34. Fitzgibbon SP, DeLosAngeles D, Lewis TW, Powers DMW, Grummett TS, Whitham EM, et al. Automatic determination of EMG-contaminated components and validation of independent component analysis using EEG during pharmacologic paralysis. Clin Neurophysiol. 2016 Mar;127(3):1781–93.

35. Rogasch NC, Sullivan C, Thomson RH, Rose NS, Bailey NW, Fitzgerald PB, et al. Analysing concurrent transcranial magnetic stimulation and electroencephalographic data: A review and introduction to the open-source TESA software. NeuroImage. 2017 Feb;147:934–51.

36. Thammasan N, Miyakoshi M. Cross-Frequency Power-Power Coupling Analysis: A Useful Cross-Frequency Measure to Classify ICA-Decomposed EEG. Sensors. 2020 Dec 9;20(24):7040.

37. Tomé D, Barbosa F, Nowak K, Marques-Teixeira J. The development of the N1 and N2 components in auditory oddball paradigms: a systematic review with narrative analysis and suggested normative values. J Neural Transm. 2015 Mar;122(3):375–91.

38. Boebinger S, Payne A, Martino G, Kerr K, Mirdamadi J, McKay JL, et al. Precise cortical contributions to sensorimotor feedback control during reactive balance. Blohm G, editor. PLOS Comput Biol. 2024 Apr 17;20(4):e1011562.

39. Taube W, Gruber M, Beck S, Faist M, Gollhofer A, Schubert M. Cortical and spinal adaptations induced by balance training: correlation between stance stability and corticospinal activation. Acta Physiol. 2007 Apr;189(4):347–58.

40. Payne AM, Ting LH. Worse balance is associated with larger perturbation-evoked cortical responses in healthy young adults. Gait Posture. 2020 July 1;80:324–30.

41. Varghese JP, Staines WR, McIlroy WE. Activity in Functional Cortical Networks Temporally Associated with Postural Instability. Neuroscience. 2019 Mar;401:43–58.

42. Payne AM, Ting LH, Hajcak G. The balance N1 and the ERN correlate in amplitude across individuals in small samples of younger and older adults. Exp Brain Res. 2023 Oct;241(10):2419–31.

43. Yazawa S, Ikeda A, Kunieda T, Mima T, Nagamine T, Ohara S, et al. Human supplementary motor area is active in preparation for both voluntary muscle relaxation and contraction: subdural recording of Bereitschaftspotential. Neurosci Lett. 1998 Mar;244(3):145–8.

44. Schurger A, Hu P “Ben”, Pak J, Roskies AL. What Is the Readiness Potential? Trends Cogn Sci. 2021 July;25(7):558–70.

45. Shibasaki H, Hallett M. What is the Bereitschaftspotential? Clin Neurophysiol. 2006 Nov;117(11):2341–56.

46. Cunnington R, Windischberger C, Robinson S, Moser E. The selection of intended actions and the observation of others’ actions: A time-resolved fMRI study. NeuroImage. 2006 Feb;29(4):1294–302.

47. Van Vugt MK, Simen P, Nystrom L, Holmes P, Cohen JD. Lateralized Readiness Potentials Reveal Properties of a Neural Mechanism for Implementing a Decision Threshold. De Lange FP, editor. PLoS ONE. 2014 Mar 13;9(3):e90943.

48. Reuter-Lorenz PA, Cappell KA. Neurocognitive Aging and the Compensation Hypothesis. Curr Dir Psychol Sci. 2008 June;17(3):177–82.

49. Reuter-Lorenz PA, Park DC. How Does it STAC Up? Revisiting the Scaffolding Theory of Aging and Cognition. Neuropsychol Rev. 2014 Sept;24(3):355–70.

